# A continuous dynamic genome-scale model explains batch fermentations led by species of the *Saccharomyces* genus

**DOI:** 10.1101/2024.05.03.592398

**Authors:** Artai R. Moimenta, Diego Troitiño-Jordedo, David Henriques, Alba Contreras-Ruíz, Romain Minebois, Miguel Morard, Eladio Barrio, Amparo Querol, Eva Balsa–Canto

## Abstract

Batch fermentation is a biotechnological dynamic process that produces various products by employing microorganisms that undergo different growth phases: lag, exponential, growth-non-growth, stationary, and decay. Genome-scale constrained-based models are commonly used to explore the phenotypic potential of these microorganisms.

Previous studies have primarily used dynamic Flux Balance Analysis (dFBA) to elucidate the metabolism during the exponential phase. However, this approach falls short in addressing the multi-phase nature of the process and secondary metabolism, posing significant challenges to our understanding of batch fermentation. A recent attempt at a solution was a discontinuous, multi-phase, multi-objective dFBA implementation.

However, this approximation lacks the mechanistic connection between phases, limiting its applicability in predicting intracellular fluxes during batch fermentation.

To overcome these limitations, we combined a novel continuous model with a genome-scale model to predict the distribution of intracellular fluxes throughout the batch fermentation process. The proposed model includes empirical descriptions of regulation that automatically identify the transition between phases. Its application to explain primary and secondary metabolism of *Saccharomyces* species in batch fermentation results in biological insights that are in good agreement with the previous literature. The ability to account for all process phases and explain secondary metabolism makes this model a valuable and easy-to-use tool for exploring novel fermentation processes.

**IMPORTANCE:** This research proposes a novel dynamic genome-scale modelling approach for batch fermentation, a crucial process widely used to produce a diverse range of products such as biofuels, enzymes, pharmaceuticals, and food products or ingredients.

The proposed approach automatically accounts for the transitions between different phases of the fermentation process (lag, exponential, growth-no-growth, and stationary). This is a significant advancement over previous methods that required different model formulations for different phases.

We have successfully applied this modelling approach to explore the primary and secondary metabolism of three yeast species under batch fermentation conditions. The model accurately explained experimental data and provided biological insights consistent with previous research findings, instilling confidence in its reliability and accuracy.

The ability of this modelling approach to explain primary and secondary metabolism makes it a valuable tool for designing novel, more efficient, and effective fermentation processes, which could have far-reaching implications in industrial biotechnology.

## Introduction

Fermentation is a commonly used technique in industrial and food biotechnology for producing a variety of products, such as antibiotics, enzymes, and biofuels. It is also the most widely used process for making fermented foods and beverages like wine, yoghurt, bread, and beer [Dimidi et al., 2019].

In most cases, fermentation runs as a dynamic batch process that occurs in several phases, with each phase playing a specific role in producing the desired product characteristics. These phases include the lag, exponential growth, growth-to-non-growth transition, stationary, and decay phases. The duration of each phase depends on factors such as the species or strain of microorganisms being used, the specific medium, and environmental conditions (e.g. temperature, pH)

Genome-scale constrained-based models (GEM) are commonly used to explore the phenotypic potential of the associated microorganism [Price et al., 2004, Palsson, 2015, Chen et al., 2022]. GEM impose environmental constraints using mass and energy balances, thermodynamics, and flux capacities. These constraints characterise all possible phenotypes of the microorganism. Flux Balance Analysis can then determine the intracellular flux distribution corresponding to a particular biological objective such as maximum biomass growth or ATP production [Orth et al., 2010].

Dynamic implementations of flux balance analysis (dFBA, Mahadevan et al. [2002]) have been successfully used to explain primary metabolism in batch fermentation. So far, most contributions reasonably explained the measured dynamics of biomass growth, carbon source uptake, and the production of relevant primary metabolites in alcoholic fermentation by *Saccharomyces cerevisiae* [Hjersted and Henson, 2009, Sánchez et al., 2014, Vargas et al., 2011, Saitua et al., 2017]. However, the multi-phase nature of batch fermentation and the fact that secondary metabolism is particularly relevant for generating flavours or aromas require alternative implementations that cope with these difficulties.

A recent contribution by Henriques et al. [2021] proposed a multi-phase, multi-objective dynamic FBA implementation that successfully explained secondary metabolism and brought novel insights into how cold-tolerant yeast species achieve redox balance. The model has been applied to explain the phenotypic differences of various yeast species in batch fermentation [Scott et al., 2023, Henriques et al., 2023].

However, this modelling approach lacked a mechanistic connection between the duration of the phases, which was otherwise estimated through data fitting. This limitation restricts the generality of the model. Additionally, the multi-phase implementation required modifying the constraints and objective functions in every other phase, complicating the implementation and application for non-experts.

To overcome these limitations, we propose combining a continuous extracellular model that can automatically describe the different phases of batch fermentation with a dynamic genome-scale model to predict the distribution of intracellular fluxes. To do so, we extend the model proposed by Moimenta et al. [2023] to predict secondary metabolism during batch fermentation, incorporating the role of transport of amino acids from the extracellular medium into the cell. The model incorporates a regulatory mechanism that can automatically detect fermentation phases. The resulting dynamic model can constrain an FBA implementation, so biomass dynamics is also posed as a constraint. At the same time, the cellular objective corresponds to a time-varying compromise between ATP and protein production.

We have applied this continuous multi-phase modelling approach to explain the metabolism of three relevant *Saccharomyces* species for the food biotechnological industry [Pérez-Torrado et al., 2018]: *Saccharomyces cerevisiae* EC1118, *Saccharomyces uvarum* BMV58, and *Saccharomyces kudriavzevii* CR85 in batch fermentation in a synthetic medium. The quality of the dynamic model compares well with the discontinuous model, with the further advantage of automatically estimating phase transitions. Notably, the dynamic metabolic flux profiles are consistent with previous observations about the different redox balance strategies used by cryotolerant species [Henriques et al., 2021, 2023].

Overall, this model represents a significant advancement in the dynamic simulation of batch fermentation, automatically accounting for all process phases and secondary metabolism. It is a versatile tool that can be easily adapted to other species or strains and used to explore novel fermentations with alternative yeast species or to engineer media or species for customised products.

## Results

### Continuous multi-phase fermentation model

In this study, we formulated a model that builds on a previous one by Moimenta et al. [2023]. The model describes the role of amino acids and introduces several adaptations to account for experimental observations and to automatically transition between the different batch fermentation phases (lag, exponential, growth-non-growth and stationary phase). Detailed model equations can be found in the Materials and Methods section. Here, we describe the model key mechanisms that are included in the final, most parsimonious model:

- The biomass dynamics is explained considering its composition in terms of carbohydrates, protein and RNA. There is no decay or transport inhibition due to ethanol.
- The lag phase occurs immediately after inoculation and persists following the Baranyi and Roberts model [Baranyi and Roberts, 1994]. The lag term affects biomass growth and assimilable nitrogen uptake.
- During fermentation, yeasts uptake assimilable nitrogen and sugars, which are used to produce biomass.
- Assimilable nitrogen comprises amino acids and ammonium, corresponding to the limiting substrate. We modelled the transport of nitrogen sources using a generalised mass action model. The corresponding growth rate follows the Monod model.
- The transport of glucose and fructose, used for fermentation, is described using a Michaelis-Menten model. Notably, our model accounts for the observation that the glucose and fructose uptake rates decrease continuously during the nitrogen starvation phase, in agreement with previous studies [Schulze et al., 1996].
- Once assimilable nitrogen is depleted, cells decelerate growth and transition to the stationary phase. During this period, cells begin to accumulate carbohydrates in the cytoplasm, leading to an increase in cellular dry weight. This accumulation is considered in a secondary growth term in the biomass equation following the model proposed by Henriques and Balsa-Canto [2021].
- As cells approach the stationary phase, due to the depletion of nitrogen and sugars, cells suffer physiological, biochemical and morphological changes. In the model, we incorporated an additional state to model global regulation of this transition.
- During alcoholic fermentation, glucose and fructose are metabolized through the glycolytic pathway into several metabolites such as ethanol, succinate, acetate, glycerol, lactate, succinate or 2-3 butanediol. We also modelled the production of acetic esters (ethyl acetate, isoamyl acetate and phenyl ethyl acetate) and higher alcohols (isobutanol, isoamyl alcohol and 2-phenyl ethanol). To explain the experimental data, we used four different production models, depending on the phase in which these compounds were released into the medium. The production rate was either proportional to the uptake of sugars (ethanol), gradually reduced over time following regulation (glycerol, acetate, lactate), delayed until nitrogen depletion (succinate, ethyl acetate, isoamyl acetate, isobutanol) or delayed until nitrogen depletion and slowly repressed during the stationary phase (ethyl acetate, phenyl acetate, 2-phenyl ethanol, 2-3 butanediol).

### The continuous and discontinuous models are equivalent in explaining external metabolites

The continuous dynamic model consists of 41 ordinary differential equations that depend on 46 unknown parameters. The model was calibrated using time series data from batch fermentations of substrates such as nitrogen sources, glucose, and fructose and products such as ethanol, glycerol, acetate, succinate, higher alcohols, and acetic acid esters.

To validate the model, we first compared its quality to recover the external dynamics and internal fluxes, comparing directly to the results reported for the discontinuous model in Henriques et al. [2021]. Note that, in that work, an ordinary differential equation described the dynamics of sucrose present in the culture medium; we have also incorporated such an equation in the continuous model for the purpose of comparison. Additionally, the dynamics of acetate and succinate were adapted to account for the consumption observed on those data. The continuous model successfully captured the process dynamics for all observables, except for those with a low signal-to-noise ratio (namely, 1-hexanol, lysine, and cysteine) for which the discontinuous model also failed. The optimal parameter values and the R-squared value of goodness of fit for all the observables can be found in Supplementary Table S1.1. The mean, median, and variance of the R-squared values of both models are similar, with a slightly better mean value for the continuous model (R^2^=0.93 *versus* R^2^=0.91). To further compare both models, we performed the Kolmogorov-Smirnov test using the corresponding R-squared values of both models to contrast their statistical distributions. Results revealed no statistical evidence that both R-squared value distributions differed at any reasonable confidence level (p-value≈0.72). Therefore, it can be concluded that both models are equivalent in goodness of fit.

### The continuous model successfully described batch fermentations led by three different species

We calibrated the model for single fermentations led by three species at 20ºC: *S. cerevisiae* EC1118, *S. uvarum* BMV58, and *S. kudriavzevii* CR85, regarded as SC-EC1118, SU-BMV58 and SK-CR85 from now on. We computed the R-squared value of goodness of fit for each observable and each strain, and the mean and median were above R^2^=0.936 and R^2^=0.98, respectively, for all strains. All values are reported in the Supplementary Table S1.2. The experimental data *versus* the corresponding model predictions are presented in Figure (**1**-A); the optimal parameters and the model versus time series data for all observables can be found in the Supplementary File.

**FIG 1.**
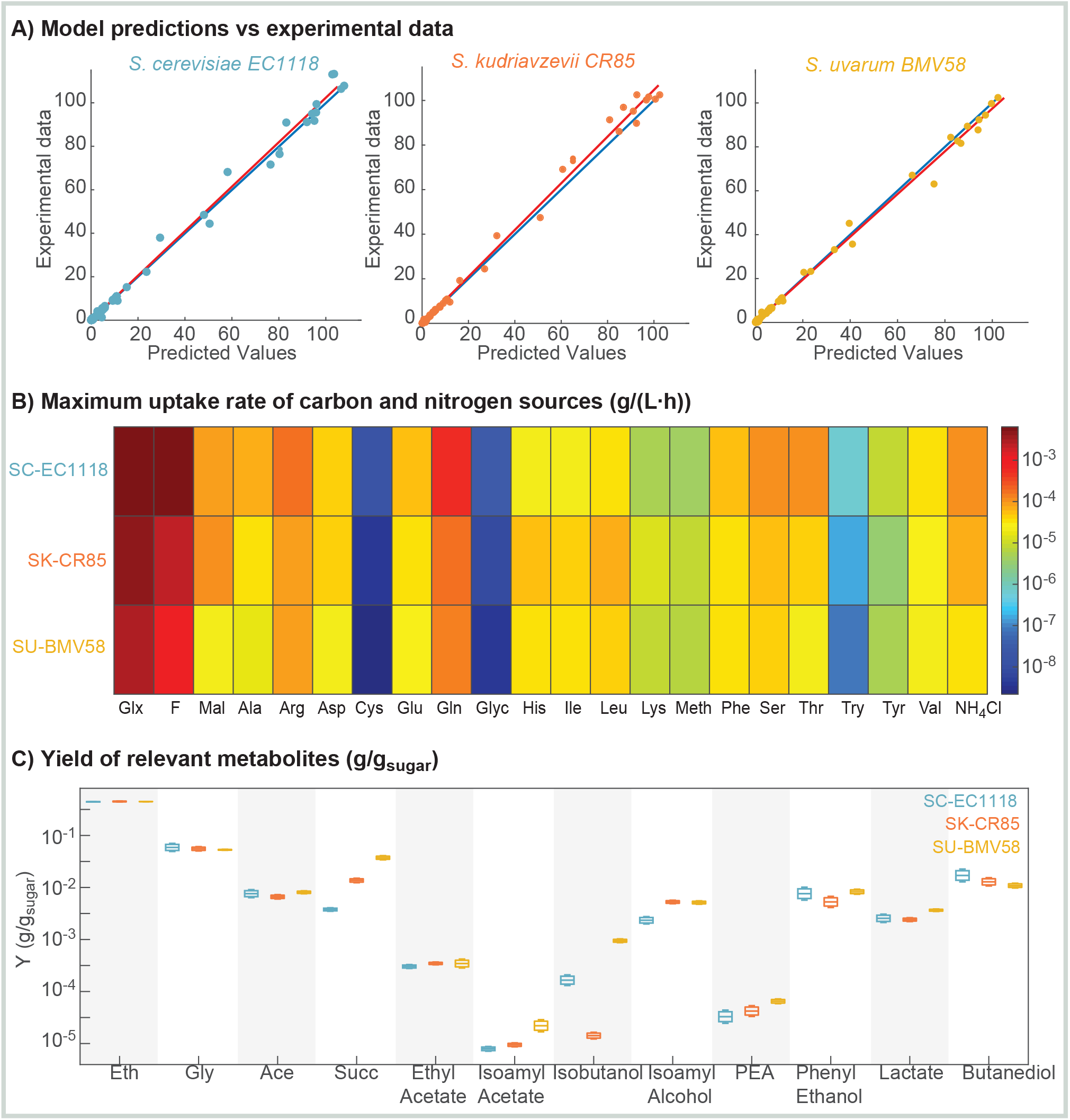
Continuous model performance. A) Experimental data *versus* model predictions for all observables. The red line shows the linear regression model to the data compared with the 1:1 line (blue line). The model recovered all data successfully with mean R^2^ ≥ 0.936 for all strains. B) Maximum uptake rate of carbon and nitrogen sources in g/(L · h). The differences show the yeast’s preference for a particular compound. C) Yields obtained for relevant metabolic products with their associated uncertainties. Substantial differences are observed in the yields of succinate, isoamyl acetate, isoamyl alcohol, phenyl ethyl acetate (PEA) or isobutanol for the three species.

Figure **1**-B shows the maximum uptake rate of carbon and nitrogen sources in g/(L · h) predicted by the model. Considerable differences are appreciated in the case of carbon source uptake among all species. SC-EC1118 seems to have more preference for both glucose and fructose (uptake rates: 6.4 *×* 10^−3^ g/(L · h) and 5.6 *×* 10^− 3^ g/(L · h), respectively) than the other species, glucose being the preferential carbon source for these three species. Particularly, SU-BMV58 presents the lowest hexoses uptake rates (2.9 *×* 10^−3^ g/(L · h) and 9.6 *×* 10^−4^ g/(L · h)). Nitrogen sources are uptaken at different rates by different species. However, cysteine and glycine are the least preferential in all cases with uptake rates between 2.2 *×* 10^−9^ g/(L · h) and 1.6 *×* 10^−8^ g/(L · h) for all species. Instead, glutamine and arginine are the most preferential, with uptake rates between 8.3 *×* 10^−5^ g/(L · h) and 5.6 *×* 10^−4^ g/(L · h). SC-EC1118 is the most efficient in uptaking nitrogen sources, while SU-BMV58 is the least efficient.

Figure **1**-C presents the yields for the relevant metabolic products and their associated uncertainties as predicted by the model. Remarkably, while the yields of ethanol, glycerol or acetate are barely distinguishable, considering the uncertainty, substantial differences appear in the production of other compounds. This is the case, for example, of succinate. Cold-tolerant species tend to produce more succinate, up to 3.6 times the yield in the case of SU-BMV58; also, a higher yield of aromatic compounds is observed for these species. For example, cold-tolerant species show twice the yield of isoamyl alcohol compared to SC-EC118.

In the sequel, we show how we used the dynamic flux balance analysis approach to explain these differences at the level of metabolic pathways.

### The metabolic reconstructions

Here, we used the Yeast8 consensus genome-scale reconstruction of *S. cerevisiae* S288C (v.8.7.0) [Lu et al., 2019] as a basis. A comparative analysis of the genome sequences of SU-BMV58 and SK-CR85 with the reference strain *S. cerevisiae* S288c was performed in previous works [Henriques et al., 2021, 2023]. In this work, we compared *S*.*cerevisiae* S288c and *S*.*cerevisiae*. SC-EC1118 was sequenced, *de novo* assembled and finally annotated with YGAP [Proux-Wéra et al., 2012]. We performed a presence/absence of genes analysis based on the functional annotation provided by YGAP. In brief, a gene from the reference strain is considered present in SC-EC1118 if YGAP gave it a systematic name. If this was not the case, then this was considered a “new” gene (present in EC1118 but not in S288C). The new genes were functionally annotated with blastKoala [Kanehisa et al., 2016]. As no relevant new or loss of functional genes was observed in the EC1118 strain genome, compared to what is considered in the reference model (see details in Supplementary Table S2.1), we used the consensus Yeast8 metabolic reconstruction.

### Automatic multi-phase-multi-objective dynamic flux balance analysis framework

We used the continuous model (9)–(25) to constrain the exchange reactions of the genome–scale model as follows:

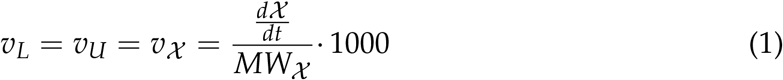

where *v*_*χ*_ represents the flux of a certain compound χ (i.e., carbon sources, nitrogen sources and all products included in the dynamic model) expressed in mmol/( DW · h^−1^). *MW χ* regards the metabolite molecular weight. However, we allowed for an inequality in the lower bound for compounds whose measurements present higher relative experimental noise. This is the case of histidine, succinate, lactate, malate, isobutanol, isoamyl alcohol, 2–phenyl acetate and 2–phenyl ethanol.

The biomass flux constraint reads as follows: (5)–(8) as

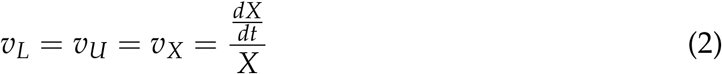

We modelled the maintenance of ATP associated with growth (*GAM*), taking into account the energetic polymerization costs of the different macromolecules, i.e. proteins, nucleic acids and carbohydrates, as:

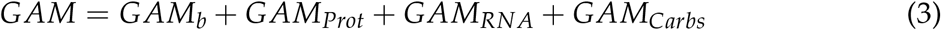

where *GAM*_*b*_ is a modifiable parameter quantifying ATP costs in growth for non-polymerisation cellular processes. In our case, we adopted the value of 30 *mmol* · *gDW*^−1^ in agreement with the previous literature [Grigaitis et al., 2023]. The model provides essential information for identifying the different fermentation phases. The lag phase may be identified during the required time when growth does not reach a significant rate. Subsequently, the primary growth phase lasts until nitrogen sources are depleted, leading to secondary growth. The metabolic transition follows *phi*_*N*_, *S* (Eqn. 24). Finally, the stationary phase is reached when the cell growth rate turns numerically null.

We used a parsimonious flux balance analysis implementation in which the cellular objective varies with time, starting with ATP maximization and ending with optimising the protein accumulation. This dynamics occurs gradually inside the cell during the different phases as follows:

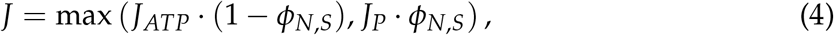

Since regulatory mechanisms, gene expression, and enzyme activity change over time, metabolic fluxes adjust to meet the evolving cellular demands. Cells prioritise ATP production in the early stages of fermentation (lag and exponential phases) to provide sufficient energy for anabolic reactions related to growth and maintenance. As cells progress through growth phases, we assume they gradually transition to maximise protein synthesis. Proteins are crucial for cell structure, enzymatic reactions, signalling, regulation and nitrogen balance. By optimising protein accumulation, cells ensure proper functioning and adaptation.

### The dynamic genome-scale modelling finds metabolic differences among species

The estimated fluxes by the model reveal important differences between strains at the extracellular level. We applied the continuous multi-phase dynamic flux balance analysis framework to decipher the most parsimonious/optimum intracellular metabolic fluxes and pathways behind these extracellular phenotypes in SC-EC1118, SU-BMV58 and SK-CR85. The simulated dynamic flux ratios are reported in Supplementary Tables S2.2-4 as mmol of the produced metabolite per mmol of consumed hexose × 100.

To analyse the differences among strains, we considered the reactions with a difference in the flux ratios given by log_10_ (|*S*_1_/*S*_2_|) ≥ 10^−3^, where *S*_1_ and *S*_2_ represent the flux ratio of a reaction for two different strains (see Supplementary Table S2.4). Figures **2** and **3** show the relevant differences. Many of the differences were found at the level of central carbon metabolism and the production of higher alcohols in the stationary phase.

**FIG 2.**
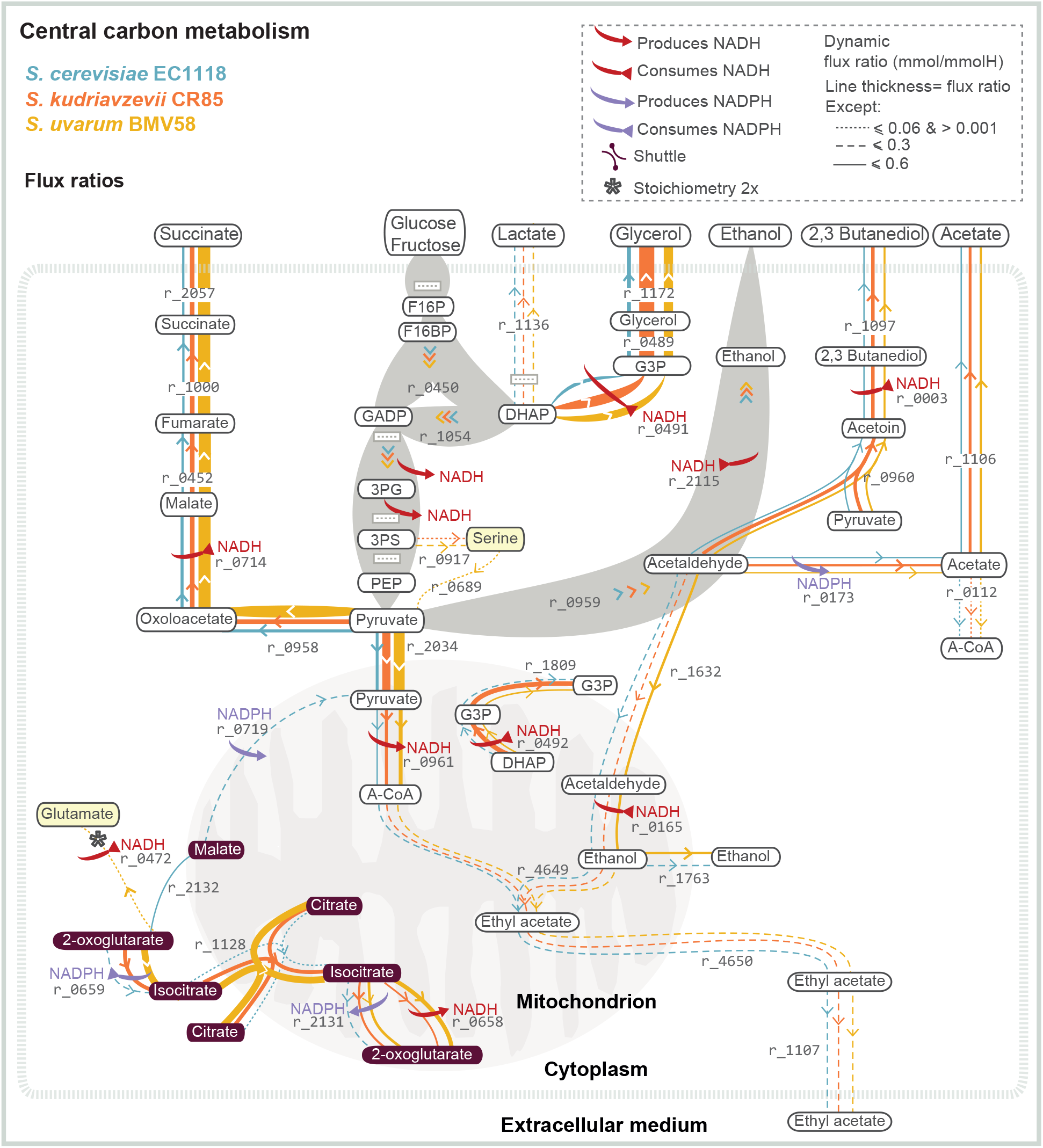
Redox balance in Central Carbon Metabolism: Figure shows the predicted intracellular dynamic flux ratios (≥ 0.01 *mmol*/*mmol H*) during the stationary phase.

**FIG 3.**
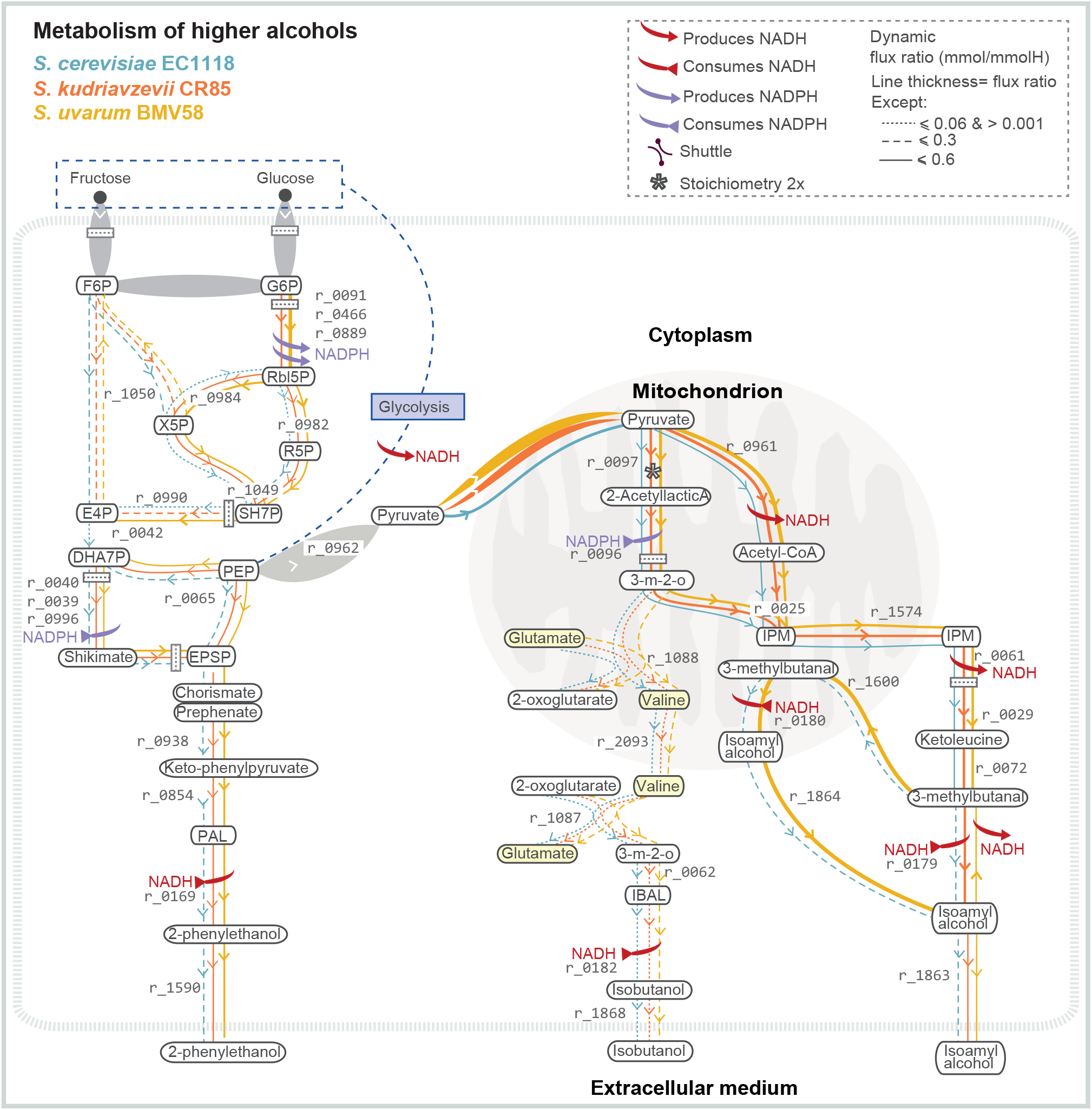
Redox balance in higher alcohol production: Figure shows the predicted intracellular flux ratios (≥ 0.01 *mmol*/*mmol H*) related to the production of higher alcohols 2–phenyl ethanol, isobutanol and isoamyl alcohol during the stationary phase as well as the role different pathways lead to the redox balance for the different species.

Although it is not the aim of the present work to explore the differences between the metabolism of the three strains in detail, here we emphasize three model outcomes:

1. **In the stationary phase, the net flux to mitochondria is substantially lower for SC-EC1118 than for SK-CR85 and SU-BMV58**. While pyruvate transport to mitochondria is higher for SC-EC1118 if we consider the flux ratios throughout the fermentation, in the stationary phase, the flux ratio is almost three and four times lower than for SK-CR85 and SU-BMV58, respectively (*r*_2034 : 1.24, 3.62, 4.87; Figure **2**). These differences are also observed in the citrate transport (*r*_1128 : 0.20, 1.70, 3.01; Figure **2**), and the production and transport of ethanol (*r*_1763 : 0.24, 3.90 *×* 10^−4^, 1.01; Figure **2**). The net flux to and from mitochondria is generally higher for SU-BMV58 in the stationary phase.
2. **Different species use different redox balance strategies**. In the case of SU-BMV58, part of the NADH available in the cytoplasm is used to produce malate from oxaloacetate (*r*_0714 : 1.32, 2.03, 5.38; Figure **2**). This would explain the significantly higher production of succinate by this species (*r*_2057 : 1.00, 2.03, 5.38; Figure **2**). SK-CR85 and SU-BMV58 use the pentose phosphate pathway differently to produce NADPH (*r*_0466, *r*_0889 : 0, 0.60, 1.59; Figure **3**), which is eventually used in the shikimate pathway (*r*_0996 : 0.18, 0.47, 0.48, Figure **3**). SU-BMV58 uses NADH produced by the action of pyruvate dehydrogenase (*r*_0961) to produce isoamyl alcohol in the mitochondria (*r*_0180 : 0.01, −7.00 *×* 10^−5^, 1.71, Figure **3**) while SK-CR85 produces isoamyl alcohol in the cytoplasm.
3. **SK-CR85 and SU-BMV58 produce higher alcohols also in the stationary phase**. Cold tolerant species show higher flux ratios of 2-phenylethanol (*r*_1590 : 0.18, 0.47, 0.48, Figure **3**) and isoamyl alcohol (*r*_1863 : 0.48, 1.05, 1.05, Figure **3**). 2-phenylethanol is produced by aldehyde dehydrogenase (ALDH), which uses NADH to reduce phenylacetaldehyde to 2-phenylethanol (*r*_0169). Isoamyl alcohol and 3-methylbutanal are produced by ketoleucine decarboxylation with NADH uptake by SC-EC1118 and SK-CR85 and with NADH production by SU-BMV58 (*r*_0179 : 0.47, 1.05, −0.66). Furthermore, although in very limited amounts, all strains produced isobutanol from valine degradation via the Ehrlich pathway in the cytoplasm (*r*_1087, *r*_0062, *r*_0182). However, these species use different routes of higher alcohol production to balance NADH.

## Discussion

Prior studies have noted the importance of using context-specific genome-scale models to improve our understanding of the link between the genome, the environment, and the phenotype of yeast species and how these models can be used to design novel bioprocesses [Chen et al., 2022]. When it comes to dynamic processes, such as batch fermentation processes, dynamic flux balance analysis (dFBA) approaches can be implemented on the basis of the quasi-steady state hypothesis [Mahadevan et al., 2002].

In a recent study by Henriques et al. [2021], a first-candidate model was proposed to explain primary and secondary metabolism in yeast batch fermentation using a multi-phase and multi-objective dFBA scheme. It successfully explained the batch fermentation process for various yeast species. Still, it lacked a mechanistic connection between the phases, which were treated as separate entities with different constraints and objectives.

In this study, we proposed a continuous extracellular dynamic model capable of detecting and explaining the different yeast batch fermentation phases. The model accounts for both primary and secondary metabolism automatically. We also combined it with the Yeast8 yeast metabolic reconstruction in a dynamic Flux Balance Analysis (dFBA) scheme. The dFBA is implemented using the continuous dynamic model to constrain external fluxes; cells are assumed to change their cellular objective throughout time, from ATP to protein maximisation. By doing so, the model predicts the dynamics of the internal metabolic flux compatible with the experimental data for biomass and extracellular compounds.

Our study builds on a previous model proposed by Moimenta et al. [2023] and suggests a continuous model that links different fermentation phases through regulation. Compared to the discontinuous model for the same species and data [Henriques et al., 2021], results are in good agreement, and the continuous model offers several computational advantages in parameter estimation and simulation. The durations of the phases are no longer parameters to be fitted to the data, improving the identifiability of the model. Furthermore, the constraints and the objective function for dFBA are unique throughout the process, simplifying the use of the model and reducing the simulation cost.

We applied the new model to explain the metabolism of three different *Saccharomyces* yeast species under batch fermentation conditions. Specifically, we selected one *S. cerevisiae* strain and two strains from the cold-tolerant species *S. uvarum* and *S. kudriavzevii*. The continuous dynamic model showed clear differences between species in the uptake rates of carbon and nitrogen sources. Consistent with the literature, the estimated glucose uptake rates were higher than that of fructose for the three species, with the *S. cerevisiae* strain consuming glucose faster than cold-tolerant species [Berthels et al., 2004, Piškur et al., 2006, López-Malo et al., 2013, Tronchoni et al., 2009]. Regarding nitrogen substrates, we reported the highest uptake rates for glutamine and arginine for the three strains. This is consistent with the position of these two amino acids as major sources and donors of nitrogen in grape must, and with the general classification of amino acids according to their quality for yeast growth [Magasanik, 2003, Crépin et al., 2012, Ljungdahl and Daignan-Fornier, 2012]. Similarly, the model showed clear differences in the yields of succinate or isoamyl alcohol, also widely documented in the literature [Minebois et al., 2020a,b, Coral-Medina et al., 2022, Pérez et al., 2022].

Our study confirms that the phenotypic differences observed in cold-tolerant species can be explained by the use of different redox balance strategies, which is consistent with our earlier observations on *S. uvarum* and *S. kudriavzevii* [Henriques et al., 2021, 2023, Minebois et al., 2020a]. However, in contrast to our previous works, the model did not identify the GABA shunt as the major metabolic pathway involved in the increase in succinate production during the stationary phase. Instead, most succinate would be produced through the TCA reductive branch. In addition to yielding succinate as an end product, the GABA shunt also produces the cofactor NADPH. In this regard, Henriques et al. [2021, 2023] proposed that the higher flux through the GABA shunt, which explained the increased succinate synthesis in SU and SK, was mainly a consequence of the need to obtain the NADPH equivalents required for lipid biosynthesis, through the consumption of acetate and the subsequent production of mevalonate (a key lipid precursor) from the start of the stationary phase. Remarkably, we did not observe acetate uptake here. One possible explanation is the fermentation medium: natural grape must vs synthetic medium used in the present work. It is plausible that the natural grape must used by Henriques et al. [2021, 2023] had a lower or even limiting lipid content, unlike the concentration used in our synthetic must (1.5 mg/L beta sitosterol; Girardi-Piva et al. [2022]). Under our non-limiting lipid conditions, *de novo synthesis* of lipids from acetate and mevalonate, as suggested by Henriques et al. [2021], has a higher energy cost than direct uptake of exogenous intermediates, located upstream in the metabolic pathway. As a result, flux through the GABA shunt becomes secondary, or even unnecessary. This hypothesis coincides in particular with the low expression of the glutamate decarboxylase encoded by GAD1 - the first step in the GABA shunt - observed by Bach et al. [2009] in synthetic grape must with non-limiting anaerobic factors (7.5 mg/L ergosterol, 2.5 mg/L oleic acid, and 0.21 g/L TWEEN80). From a modelling point of view, another possible explanation is that phase durations are slightly different when estimated through data fitting and when using the continuous model. Indeed, when estimated as parameters in the discontinuous model, their values had an associated uncertainty of up to 20%.

Therefore, the use of the GABA shunt in the different phases of the fermentation requires further analyses. The model also showed a substantial flux toward the mitochondria in the case of *S. uvarum* and larger production of higher alcohols during the stationary phase for both *S. kudriavzevii* and *S. uvarum* as compared to *S. cerevisiae*. Again, these results are in good agreement with previous findings [Henriques et al., 2021, 2023, Minebois et al., 2020a]. Note that higher alcohol production contributes to achieving the cellular redox balance for these species. Importantly, the estimated fluxes of pyruvate to the synthesis of aromas in the mitochondria support the observation by Crépin et al. [2017] that higher alcohols and their esters are from newly synthesized precursors rather than from the catabolism of exogenous amino acids.

The fact that internal fluxes are consistent with previous modelling and experimental results leads us to conclude that the proposed continuous dFBA approach allows us to link yeast genotypes with context-specific phenotypes in batch fermentation conditions. The model automatically accounts for phase transitions and uses a single formulation of constraints and objective functions within the FBA approach, facilitating its use. By considering all process phases, our model provides an accurate explanation of primary and secondary metabolism and an improved understanding of the underlying mechanisms of batch fermentation.

## Materials and Methods

### Experimental methods

#### Yeast strains

In this study, three yeast strains were utilized, specifically strains belonging to the species *S. cerevisiae, S. uvarum* and *S. kudriavzevii*. The three strains used were *S. cerevisiae* EC1118 (denoted as SC-EC118), *S. uvarum* BMV58 (denoted as SU-BMV58), and *S. kudriavzevii* CR85 (denoted as SK-CR85). SC-EC1118 is a commercial wine strain that was originally isolated from wine production in France. SU-BMV58 is another commercial wine strain, while SK-CR85 is a natural isolate obtained from an oak tree’s bark. The strains were cryogenically preserved at -80ºC and cultured and maintained on GPY plates (2% glucose, 2% agar, 0.5% peptone, 0.5% yeast extract). The GPY plates were kept at 4ºC to preserve the strains.

Prior to inoculation, an overnight starter culture was prepared by pouring a small amount of biomass from the plate into an Erlnemeyer flask with 25 mL of GPY liquid medium (2% glucose, 0.5% peptone, 0.5% yeast extract) and placed at 25^*o*^ *C* with agitation under aerobic conditions.

#### Fermentations and sampling

Time series data were obtained from fermentations carried out in temperature-controlled bioreactors at 20^*o*^ *C*. Bioreactors were filled with 470 ml of synthetic must, following the same recipe as in Rollero et al. [2019], and inoculated at OD-600 0.1 (approximately 1 · 10^6^ cells/ml) from starter cultures. For each strain, three independent biological replicates were conducted.

Samples were collected across the fermentation, covering the different phases of fermentation. At each sampling point, dry weight (DW), residual hexoses, residual amino acids and ammonium, organic acids, the main fermentative by-products (e.g. ethanol, succinate, glycerol) and volatile compounds were quantified. Cell density (C/mL) was measured using a MUSE cell analyser (Millipore). Dry weight biomass was obtained by measuring the weight difference of a pre-weighted filter (0.45 *μm* pore size) used to filtrate 5 mL of broth, rinsed twice with 50 mL of deionized water and placed for 48h at 110^*o*^ *C*. Determination of amino acids and ammonia (yeast assimilable nitrogen) was carried out following the same protocol as the one followed in Su et al. [2020]. Sugars (glucose, fructose), fermentative by-products (glycerol, ethanol, 2-3 butanediol, and erythritol), and organic acids (acetate, succinate, tartrate, citrate, and malate) were determined using an HPLC (Thermo Fisher Scientific, Waltham, MA) equipped with a refraction index and UV/VIS (210nm) detector. Samples were diluted and filtered through a 0.22-*μm* nylon filter (Symta, Madrid, Spain). Metabolite separation was performed through a HyperREZTM XP Carbohydrate H+ 8 mm column fitted with a HyperREZTM XP Carbohydrate Guard (Thermo Fisher Scientific, Waltham, MA). The mobile phase consisted of 1.5 mM of H2SO4 at a flux of 0.6 ml/min, and the HPLC oven was kept a 50 ^*o*^ *C*.

For more detailed information on the experimental design and the methods of measurement of the different compounds analysed see article Contreras-Ruiz et al. (Preprint, paper in progress), since the biological data used in this article come from that previously conducted study.

### Theoretical methods

#### Dynamic continuous model

The model builds upon a previously developed model by Moimenta et al. [2023]. The model accounts for the following state variables: biomass (*X*) (g_DW_/L); protein content in biomass (*X*_*P*_) (g_P_/L); carbohydrate content in biomass (*X*_*C*_) (g_C_/L); mRNA content in biomass (*X*_*mRNA*_) (g_mRNA_/L); glucose and fructose (*Glx* and *F*); ethanol (*Eth*,g/L); glycerol (*Gly*); succinate (*Succ*); acetate (*Ace*); several secondary metabolites such as ethyl acetate (*Ethyl Acetate*), isoamyl acetate (*Isoamyl Acetate*), phenyl ethyl acetate (*PEA*), isobutanol (*Isobutanol*), isoamyl alcohol (*Isoamyl Alcohol*), 2-3 butanediol (*Butanediol*), 2-phenyl ethanol (*PE*) and malate (*Malate*). All the metabolites are measured in (g/L). The model accounts for assimilable nitrogen (*YAN*) (g_N_/L) including ammonia (*NH*4) (g_NH4_/L) and the different amino acids measured in (g/L) present in the medium: alanine (*Ala*), arginine (*Arg*), aspartate (*Asp*), cysteine (*Cys*), glutamate (*Glu*), glutamine (*Gln*), glycine (*Gly*), histidine (*His*), isoleucine (*Ile*), leucine (*Leu*), lysine (*Lys*), methionine (*Meth*), phenylalanine (*Phe*), serine (*Ser*), threonine (*Thr*), tyrosine (*Tyr*), tryptophan (*Try*) and valine (*Val*). Note that yeasts do not metabolize proline under anaerobic conditions; therefore, we do not include it as an assimilable nitrogen source.

The model was built following an iterative procedure. Here we present the original, most complete model, which accounts for cellular decay:

- Differential equations:
  - Biomass production. Biomass is modelled taking into account its major components: carbohydrates, protein and mRNA. Their dynamics are described as follows:

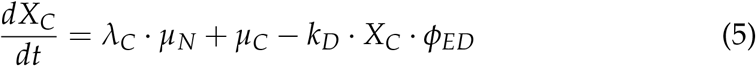

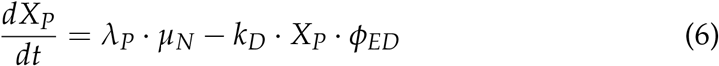

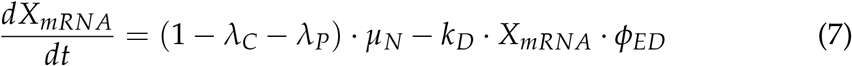

where *λ*_*C*_ = 0.29, *λ*_*P*_ = 0.59 and (1 − *λ*_*C*_ − *λ*_*P*_ = 0.12) correspond to the percentage of carbohydrate, protein and mRNA in biomass respectively. We used the values reported by Schulze et al. [1996] for nitrogen-limited batch fermentation (59% protein and 12% mRNA); *μ*_*N*_ and *μ*_*C*_ (1/*h*) regard primary and secondary growth; and *k*_*D*_ · *ϕ*_*ED*_ represent the cellular decay due to the production of ethanol. The biomass equation results:

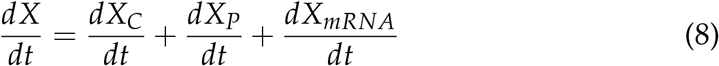
  - Nitrogen uptake.

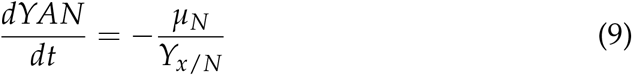

*Y*_*xN*_ (g_DW_/g_N_) is the yield of nitrogen to biomass.
  - Ammonia and amino acids uptake follows a generalised mass action model:

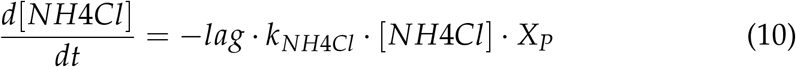

and

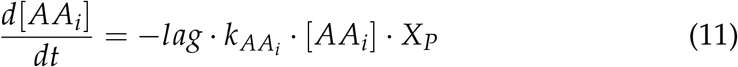

where we denoted by [*AA*_*i*_] the extracellular concentration of each amino acid in *g*/*L* and 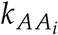 (1/(g_prot_ · h)) is the associated kinetic parameter.
  - Sugars uptake. The medium contains both glucose and fructose. The uptake is described following the Michaelis-Menten model. Note that the uptake rate varies when nitrogen is still available in the medium.

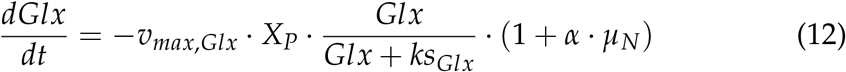

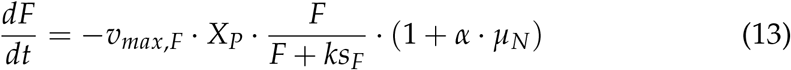

Where *v*_*max,Glx*_ and *v*_*max,F*_ correspond to the maximum uptake rates for glucose and fructose, respectively, in g/L. *ks*_*Glx*_ and *ks*_*F*_ correspond to the Michaelis-Menten constants. The term *α* · *μ*_*N*_ describes the increase in the production of sugar transporters during primary growth, which results in a higher uptake rate of sugars during that phase.
  - Gene regulation. An empirical expression describes the transition to the stationary phase once YAN and sugars are being depleted:

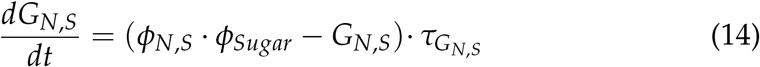

*ϕ*_*N,S*_ and *ϕ*_*Sugar*_ are smooth sigmoidal functions used to activate secondary growth and transition to the stationary phase (see their definitions later in this section). 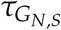 controls the velocity of those transcriptional changes.
  - Extracellular primary metabolites production. The rate of excretion of metabolites is directly proportional to the glucose and fructose uptake. The yield coefficient (*Y*_*i*_, where *i* represents Eth, Gly, Ace, Succ) expressed in (g_i_/g_sugar_) varies significantly depending on the fermentation phase. We use smooth logistic type functions to describe this variation, depending on whether there is an increase in production associated with the transition to secondary growth (*Y*_*i*_ · *ϕ*_*N,S*_), or a decrease in production associated with the transition to the stationary phase (*Y*_*i*_ · (1 − *G*_*N,S*_)).

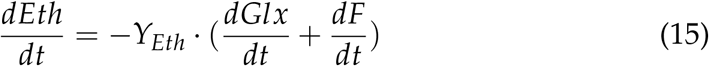

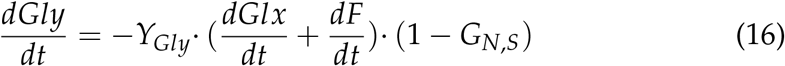

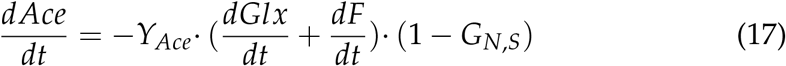

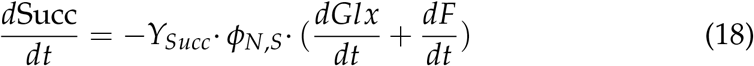
  - Extracellular secondary metabolites production. Analogous expressions are used to describe secondary metabolites. Metabolites modelled were acetic esters (Ethyl Acetate, Isoamyl Acetate and Phenyl Ethyl Acetate), higher alcohols (Isobutanol, Isoamyl Alcohol, 2-phenyl Ethanol) and 2,3 butanediol.

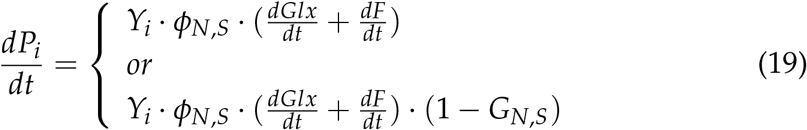

The best candidate in Equation (19) was selected for each product through model calibration.
  - Experimental data revealed that under tested conditions, cells uptake malic acid which we described following a generalised mass action model:

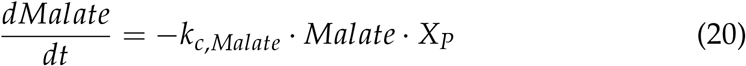

where *k*_*c,Malate*_ (1/(g_prot_ · h)) is the associated kinetic parameter.
- Constitutive equations:
  - Lag phase.

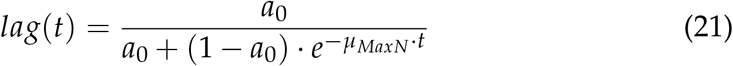

Analytical solution of the commonly accepted model by Baranyi and Roberts [1994]. *a*_0_ is a parameter bounded between 0 and 1 and represents the physiological state of the inoculum. *μ*_*maxN*_ is the maximum specific growth rate (h^−1^). This function enables the transition from the lag phase to the exponential growth phase.
  - Primary growth.

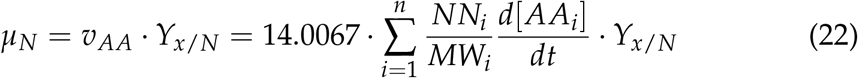

*μ*_*N*_ describes the specific growth rate associated with cell division. We defined growth considering the proportion of nitrogen present in the amino acids and ammonia. *NN*_*i*_ and *MW*_*i*_ refer to the number of nitrogen atoms and molecular weight present in the molecule of each nitrogen compound (amino acid and ammonia), respectively, e.g. *Arginine*: *NN*_*Arg*_ = 4; *MW*_*Arg*_ = 174.2g/mol.
  - Secondary growth.

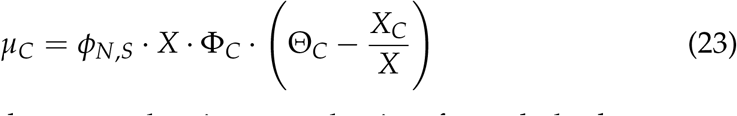

*μ*_*C*_ describes the secondary growth using a mechanism for carbohydrate accumulation which was modelled with an empirical expression analogous to a proportional controller [Henriques and Balsa-Canto, 2021]. Θ_*C*_ represents a set point for the ratio of carbohydrates in the biomass content 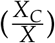 and Φ_*C*_ controls the velocity of the convergence towards that point. The activation of the secondary growth was modelled using the function (*ϕ*_*N,S*_):

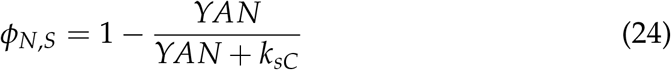

*ϕ*_*N,S*_ regulates the nitrogen concentration needed to induce carbohydrate accumulation and the transcriptional changes associated with nitrogen starvation. *k*_*sC*_ is the half-saturation constant in g/L. Note that *ϕ*_*N,S*_ corresponds to a sigmoidal function bounded between [0, 1]. Its value is close to 0 at the beginning of the process while nitrogen sources are still available and rapidly becomes 1 fully activating the secondary growth.
  - Transition to the stationary phase. Cells transition to the stationary phase once nitrogen has been depleted and sugars are gradually being uptaken. We defined a function *ϕ*_*Sugar*_, that regulates the sugar concentration needed to induce the transcriptional changes associated with the stationary phase.

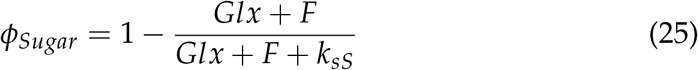

#### Model calibration

The model in Equations (5)-(25) depends on more than 40 unknown parameters, which are to be estimated from experimental data. The so-called model calibration problem aims to find the values of the unknown parameters that minimize (or maximize) some distance between model predictions and experimental data, subject to the system dynamics and parameter bounds [Vilas et al., 2017]. Hence, the optimal parameters correspond to those that maximize the (log-)likelihood function:

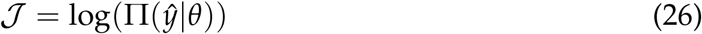

Under the assumptions of independently identically distributed measurements according to a Gaussian law for a given sampling time *t*_*i*_, the maximization of 𝒥 is equivalent to minimizing:

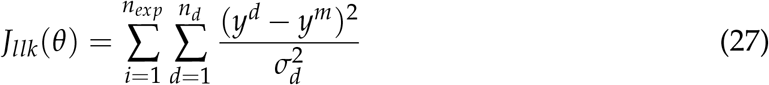

where *n*_*exp*_ corresponds to the number of experiments; *n*_*d*_ corresponds to the number of data per experiment; *y*^*m*^ and *y*^*d*^ regard the model predictions and the measured data, respectively, and *σ*_*d*_ is the standard deviation associated to the experimental data as computed from the experimental replicates.

The confidence interval (*ρ*_*i*_) associated with each parameter estimate may be obtained through the covariance matrix:

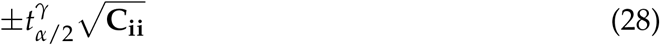

where 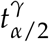 is given by Students t-distribution, *γ* = *N*_*d*_ − *η* degrees of freedom and (1 − *α*)100% is the confidence interval selected, typically 95%.

For non-linear models, there is no exact way to obtain **C**. Therefore, approximations have been suggested. Possibly the most widely used is based on the Crammèr-Rao inequality, which establishes, under certain assumptions on the number of data and non-linear character of the model, that the covariance matrix may be approximated by the inverse of the Fisher information matrix (ℱ) in its typical definition:

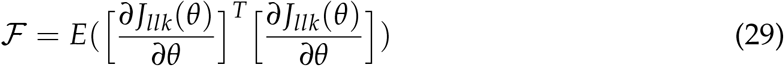

#### Parsimonious flux balance analysis

Metabolic fluxes in the cell are obtained using a constraint-based model derived from flux balance analysis (FBA) [Varma and Palsson, 1994, Orth et al., 2010]. FBA is a modelling framework based on the metabolic network stoichiometry and a steady–state mass balance condition:

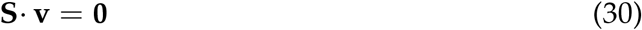

where **S** is the stoichiometric matrix of (n metabolites by m reactions) and **v** is a vector of metabolic fluxes. The number of unknown fluxes is higher than the number of equations, and thus, the system is undetermined. Still, it is possible to find a unique solution under the assumption that cell metabolism complies with the maximisation (or minimisation) of a certain cellular objective (*J*):

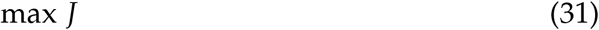

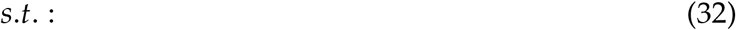

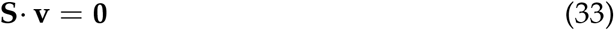

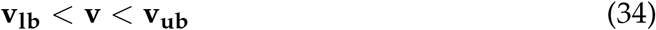

where **v**_**lb**_ and **v**_**ub**_ correspond to the lower and upper limits of the estimated fluxes. Examples of FBA problems include maximizing growth rate or ATP production, minimizing uptake of nutrients, etc.

Further, parsimonious FBA is then used to minimize the flux while maintaining optimum flux through the objective function [Machado and Herrgård, 2014].

### Analysis of the dynamic metabolic fluxes

We selected the most relevant metabolic pathways using a flux ratio, which measures the net flux throughout the growth and stationary phases. In particular, we calculated the integral of each flux multiplied by biomass over time and normalized its value with the accumulated flux of consumed hexoses:

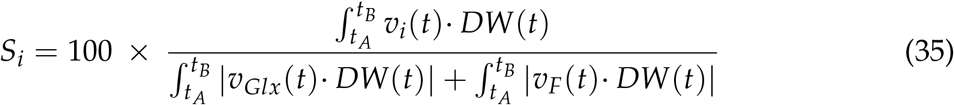

where *S*_*i*_ corresponds to the flux score *i, v*_*i*_(*t*) (mmol/(h · DW)) is the flux under scrutiny, *v*_*Glx*_(*t*) (mmol/(h · DW)) corresponds to the glucose flux, equivalently *v*_*F*_(*t*) (mmol/(h · DW)) corresponds to the fructose flux, DW is the predicted dry weight biomass (g) and *t*_*A*_ (hours) and *t*_*B*_ (hours) define the interval of interest. The results correspond to the mmol of compound produced per mmol of hexoses consumed *×*100 (denoted as mmol/mmolH).

### Numerical tools

To automatise the model calibration, we used the AMIGO2 toolbox [Balsa-Canto et al., 2016]. AMIGO2 is a MATLAB based toolbox focused on dynamic model identification and optimization. AMIGO2 includes sensitivity and identifiability analyses and offers several numerical methods for simulation and optimization. Specifically, we used CVODES [Hindmarsh et al., 2005] to solve the model and the *enhanced Scatter Search* method [eSS, Egea et al., 2009] to optimize parameters. AMIGO2 automatically reports the parametric confidence intervals as obtained by means of the Crammèr-Rao inequality.

To compare the discontinuous and continuous models, we performed a Kolmogorov–Smirnov test over the R^2^ values of both models using the ks.test included in the R package stats [Wickham, 2016].

To solve the dFBA problem, we have coupled AMIGO initial value problem solver (CVODES) with COBRA Toolbox [Schellenberger et al., 2011]. We used a variable–step, variable–order Adams–Bashforth–Moulton method to solve the initial value problem defined by the system of ordinary differential equations that describe the dynamics of the extracellular metabolites. At each time step determined by this method, a pFBA problem was performed using COBRA Toolbox.

Integrals required to compute flux ratios were evaluated using the standard trapezoidal method (function *trapz* in MATLAB).

All scripts necessary to reproduce the results are distributed in https://sites.google.com/site/amigo2toolbox/examples.

## Acknowledgments

This work has received funding from MCIU/AEI/FEDER grant references: PID2021-126380OB-C31, PID2021-126380OB-C32, PID2021-126380OB-C33 and Xunta de Galicia (IN607B 2023/04). IATA-CSIC received funding from the Spanish Government, ref. MCIN/AEI/10.13039/501100011033, as a ‘Severo Ochoa’ Center of Excellence (CEX2021-001189-S), with AQ as PI.

## References

Dimidi E, Cox SR, Rossi M, Whelan K. Fermented Foods: Definitions and Characteristics, Impact on the Gut Microbiota and Effects on Gastrointestinal Health and Disease. Nutrients 2019;11(8).

Price ND, Reed JL, Palsson BO. Genome-scale models of microbial cells: evaluating the consequences of constraints. Nat Rev Microbiol 2004;2(11):886–97.

Palsson BO. Systems Biology: Constraint-based Reconstruction and Analysis. Cambridge University Press; 2015.

Chen Y, Li F, Nielsen J. Genome-scale modeling of yeast metabolism: retrospectives and perspectives. FEMS Yeast Research 2022;22(1):foac003.

Orth JD, Thiele I, Palsson BØ. What is flux balance analysis? Nature biotechnology 2010;28(3):245–248.

Mahadevan R, Edwards JS, Doyle III FJ. Dynamic flux balance analysis of diauxic growth in Escherichia coli. Biophysical journal 2002;83(3):1331–1340.

Hjersted JL, Henson MA. Steady-state and dynamic flux balance analysis of ethanol production by Saccharomyces cerevisiae. IET Systems Biology 2009;3(3):167–179.

Sánchez BJ, Pérez-Correa JR, Agosin E. Construction of robust dynamic genome-scale metabolic model structures of Saccharomyces cerevisiae through iterative re-parameterization. Metabolic engineering 2014;25:159–173.

Vargas FA, Pizarro F, Pérez-Correa JR, Agosin E. Expanding a dynamic flux balance model of yeast fermentation to genome-scale. BMC systems biology 2011;5(1):75.

Saitua F, Torres P, Pérez-Correa JR, Agosin E. Dynamic genome-scale metabolic modeling of the yeast Pichia pastoris. BMC Systems Biology 2017;11(1).

Henriques D, Minebois R, Mendoza SN, Macías LG, Pérez-Torrado R, Barrio E, et al. A multiphase multiobjective dynamic genome-scale model shows different redox balancing among yeast species of the Saccharomyces genus in fermentation. mSystems 2021;6(4).

Scott WT, Henriques D, Smid EJ, Notebaart RA, Balsa-Canto E. Dynamic genome-scale modeling of Saccharomyces cerevisiae unravels mechanisms for ester formation during alcoholic fermentation. Biotechnology and Bioengineering 2023;120(7):1998 –2012.

Henriques D, Minebois R, dos Santos D, Barrio E, Querol A, Balsa-Canto E. A Dynamic Genome-Scale Model Identifies Metabolic Pathways Associated with Cold Tolerance in Saccharomyces kudriavzevii. Microbiology Spectrum 2023;11(3):e03519–22.

Moimenta AR, Henriques D, Minebois R, Querol A, Balsa-Canto E. Modelling the physiological status of yeast during wine fermentation enables the prediction of secondary metabolism. Microbial Biotechnology 2023;16(4):847 – 861.

Pérez-Torrado R, Barrio E, Querol A. Alternative yeasts for winemaking: Saccharomyces non-cerevisiae and its hybrids. Crit Rev Food Sci Nut 2018;58(11):1780 –1790.

Baranyi J, Roberts TA. A dynamic approach to predicting bacterial growth in food. Int J Food Microbiol 1994;23:277–294.

Schulze U, Lidén G, Nielsen J, Villadsen J. Physiological effects of nitrogen starvation in an anaerobic batch culture of Saccharomyces cerevisiae. Microbiology 1996;142(8):2299–2310.

Henriques D, Balsa-Canto E. The Monod Model Is Insufficient To Explain Biomass Growth in Nitrogen-Limited Yeast Fermentation. Appl Environ Microbiol 2021;87(20):e01084–21.

Lu H, Li F, Sánchez BJ, Zhu Z, Li G, Domenzain I, et al. A consensus S. cerevisiae metabolic model Yeast8 and its ecosystem for comprehensively probing cellular metabolism. Nature communications 2019;10(1):3586.

Proux-Wéra E, Armisén D, Byrne KP, Wolfe KH. A pipeline for automated annotation of yeast genome sequences by a conserved-synteny approach. BMC Bioinf 2012;13(1):1–12.

Kanehisa M, Sato Y, Morishima K. BlastKOALA and GhostKOALA: KEGG Tools for Functional Characterization of Genome and Metagenome Sequences. J Mol Biol 2016;428(4):726–731.

Grigaitis P, van den Bogaard SL, Teusink B. Elevated energy costs of biomass production in mitochondrial respiration-deficient Saccharomyces cerevisiae. FEMS Yeast Research 2023;23:foad008.

Berthels NJ, Cordero Otero RR, Bauer FF, Thevelein JM, Pretorius IS. Discrepancy in glucose and fructose utilisation during fermentation by Saccharomyces cerevisiae wine yeast strains. FEMS Yeast Res 2004;4(7):683–689.

Piškur J, Rozpedowska E, Polakova S, Merico A, Compagno C. How did Saccharomyces evolve to become a good brewer? Trends Genetics 2006;22(4):183 – 186.

López-Malo M, Querol A, Guillamón JM. Metabolomic Comparison of Saccharomyces cerevisiae and the Cryotolerant Species S. bayanus var. uvarum and S. kudriavzevii during Wine Fermentation at Low Temperature. PLOS ONE 2013;8(3):1–14.

Tronchoni J, Gamero A, Arroyo-Lopez N, Barrio E, Querol A. Differences in the glucose and fructose consumption profiles in diverse Saccharomyces wine species and their hybrids during grape juice fermentation. International Journal of Food Microbiology 2009;134(3):237 – 243.

Magasanik B. Ammonia assimilation by Saccharomyces cerevisiae. Eukaryot Cell 2003;2(5):827–829.

Crépin L, Nidelet T, Sanchez I, Dequin S, Camarasa C. Sequential Use of Nitrogen Compounds by Saccharomyces cerevisiae during Wine Fermentation: a Model Based on Kinetic and Regulation Characteristics of Nitrogen Permeases. Applied and Environmental Microbiology 2012;78(22):8102–8111.

Ljungdahl PO, Daignan-Fornier B. Regulation of amino acid, nucleotide, and phosphate metabolism in Saccharomyces cerevisiae. Genetics 2012;190(3):885–929.

Minebois R, Pérez-Torrado R, Querol A. A time course metabolism comparison among Saccharomyces cerevisiae, S. uvarum and S. kudriavzevii species in wine fermentation. Food Microbiol 2020;90:103484.

Minebois R, Pérez-Torrado R, Querol A. Metabolome segregation of four strains of Saccharomyces cerevisiae, Saccharomyces uvarum and Saccharomyces kudriavzevii conducted under low temperature oenological conditions. Environ Microbiol 2020;22(9):3700–3721.

Coral-Medina A, Morrissey JP, Camarasa C. The growth and metabolome of Saccharomyces uvarum in wine fermentations are strongly influenced by the route of nitrogen assimilation. J Ind Microbiol Biotechnol 2022;49(6):kuac025.

Pérez D, Denat M, Minebois R, Heras JM, Guillamón JM, Ferreira V, et al. Modulation of aroma and chemical composition of Albariño semi-synthetic wines by non-wine Saccharomyces yeasts and bottle aging. Food Microbiol 2022;104:103981.

Girardi-Piva G, Casalta E, Legras JL, Nidelet T, Pradal M, Macna F, et al. Influence of ergosterol and phytosterols on wine alcoholic fermentation with Saccharomyces cerevisiae strains. Front Microbiol 2022;13:966245.

Bach B, Sauvage FX, Dequin S, Camarasa C. Role of γ-Aminobutyric Acid as a Source of Nitrogen and Succinate in Wine. Am J Enol Vit 2009;60(4):508–516.

Crépin L, Truong NM, Bloem A, Sanchez I, Dequin S, Camarasa C. Management of multiple nitrogen sources during wine fermentation by Saccharomyces cerevisiae. Applied and environmental microbiology 2017;83(5):e02617–16.

Rollero S, Bloem A, Ortiz-Julien A, Bauer FF, Camarasa C, Divol B. A comparison of the nitrogen metabolic networks of Kluyveromyces marxianus and Saccharomyces cerevisiae. Environmental Microbiology 2019;21(11):4076–4091.

Su Y, Seguinot P, Sanchez I, Ortiz-Julien A, Heras JM, Querol A, et al. Nitrogen sources preferences of non-Saccharomyces yeasts to sustain growth and fermentation under winemaking conditions. Food Microbiology 2020;85:103287.

Vilas C, Arias-Méndez A, Garcia MR, Alonso AA, Balsa-Canto E. Towards predictive food process models: A protocol for parameter estimation. Crit Rev Food Sci & Nut 2017;.

Varma A, Palsson BO. Stoichiometric flux balance models quantitatively predict growth and metabolic by-product secretion in wild-type Escherichia coli W3110. Applied and environmental microbiology 1994;60(10):3724–3731.

Machado D, Herrgård M. Systematic evaluation of methods for integration of transcriptomic data into constraint-based models of metabolism. PLoS computational biology 2014;10(4):e1003580.

Balsa-Canto E, Henriques D, Gabor A, Banga JR. AMIGO2, a toolbox for dynamic modeling, optimization and control in systems biology. Bioinformatics 2016;32(21):3357–3359.

Hindmarsh AC, Brown PN, Grant KE, Lee SL, Serban R, Shumaker DE, et al. SUNDIALS: Suite of Nonlinear and Differential/Algebraic Equation Solvers. ACM Trans Math Softw 2005;31(3):363–396.

Egea JA, Vazquez E, Banga JR, Marti R. Improved scatter search for the global optimization of computationally expensive dynamic models. J Global Opt 2009;43(2–3):175–190.

Wickham H. ggplot2: Elegant Graphics for Data Analysis. Springer-Verlag New York; 2016.

Schellenberger J, Que R, Fleming R, Thiele I, Toolbox J, Feist A, et al. Quantitative prediction of cellular metabolism with constraint-based models: the COBRA Toolbox v2.0. Nat Protoc 2011;4(6(9)):1290–307.

